# Comparison of one-stage and two-stage genome-wide association studies

**DOI:** 10.1101/099291

**Authors:** Shang Xue, Funda Ogut, Zachary Miller, Janu Verma, Peter J. Bradbury, James B. Holland

## Abstract

Linear mixed models are widely used in humans, animals, and plants to conduct genome-wide association studies (GWAS). A characteristic of experimental designs for plants is that experimental units are typically multiple-plant plots of families or lines that are replicated across environments. This structure can present computational challenges to conducting a genome scan on raw (plot-level) data. Two-stage methods have been proposed to reduce the complexity and increase the computational speed of whole-genome scans. The first stage of the analysis fits raw data to a model including environment and line effects, but no individual marker effects. The second stage involves the whole genome scan of marker tests using summary values for each line as the dependent variable. Missing data and unbalanced experimental designs can result in biased estimates of marker association effects from two-stage analyses. In this study, we developed a weighted two-stage analysis to reduce bias and improve power of GWAS while maintaining the computational efficiency of two-stage analyses. Simulation based on real marker data of a diverse panel of maize inbred lines was used to compare power and false discovery rate of the new weighted two-stage method to single-stage and other two-stage analyses and to compare different two-stage models. In the case of severely unbalanced data, only the weighted two-stage GWAS has power and false discovery rate similar to the one-stage analysis. The weighted GWAS method has been implemented in the open-source software TASSEL.

## Background

Genome-wide association studies are widely used to identify genes affecting complex traits in humans, animals, and plants (Stange *et al.* 2013; Zhang *et al.* 2012; Huang and Han 2014). Large sample sizes are required to achieve good statistical power in GWAS (Balding 2006). In addition, the number of markers to be tested is increasing rapidly as next-generation genomics techniques permit the acquisition of dense genome-wide marker data. In addition to the high dimensionality of the data, marker tests need to be adjusted for population structure and genomic relationships (Yu *et al.* 2006). Combined, these factors result in significant computational burden for GWAS in many instances.

A common problem in GWAS is the need to account for extraneous non-genetic factors that affect the phenotypes. Estimating the effect associated with a single SNP while also including extraneous factors and modeling genetic background effects using a complex variance-covariance structure in linear mixed models can dramatically increase computation time for each marker test. To reduce computational demands for linear mixed model-based GWAS, several different strategies have been proposed in human, animal and plant studies to simplify and speed up computation time for the individual marker tests. In general, these strategies approach the problem using stage-wise procedures that first adjust the phenotypes for extraneous effects and second conduct linear mixed model GWAS on the adjusted phenotypes. For example, in human GWAS studies, two-stage regression analysis is a widely used strategy to test SNPs for associations with quantitative diseases (Laird *et al.* 2000; Zeegers *et al.* 2004; Naylor *et al.* 2009). First, residual effects (‘adjusted-outcomes’) for each individual are calculated by regressing the raw phenotype on covariates such as demographic, clinical, and environmental factors. Second, the residual values are used as the dependent variable to test the association with SNP markers using a simple linear regression. Although this approach greatly reduces computational burden, it results in biased estimates of genotypic effects and reduced power (Demissie and Cupples 2011; Che *et al.* 2012).

Similarly, GWAS of domesticated animals often uses stage-wise approaches. In animal studies, the researcher may have available raw phenotypes of individuals and also their estimated breeding values (EBV) from pedigree-based analyses of historical data. Some animals may have genotypes and EBVs based on information from their relatives, but no direct phenotypes. Although EBVs have been used as dependent variables in GWAS (Becker *et al.* 2013; Johnston *et al.* 2011), this approach has a high false positive rate (Ekine *et al.* 2013). Consequences of using EBVs include varying levels of precision and ‘shrinkage effect’ among the values used to represent phenotypes of different individuals, a reduction in the sample variance of the phenotypes, and ‘double-counting’ of information from relatives (Garrick *et al.* 2009; Ostersen *et al.* 2011). As an alternative, the EBVs can be ‘deregressed’ (Garrick *et al.* 2009; Ostersen *et al.* 2011) to standardize the variance and influence of the individuals’ EBVs while still accounting for information from relatives. The use of deregressed EBVs as dependent variables can improve the power of GWAS (Sevillano *et al.* 2015; Sell-Kubiak *et al.* 2015). Another alternative is to fit a mixed model to the data on individuals, accounting for either pedigree relationships or realized genomic relationships, then use the residuals for each individual as the dependent variable in a second stage genomic scan (Amin et al. 2007; Lam et al. 2007; Aulchenko et al. 2007).This approach,called‘GRAMMAR’, dramatically speeds up computation time for the GWAS because each marker test is a simple linear regression of residual values on genotype scores at one locus, however, this approach has low power in some cases (Zhou and Stephens 2012).

In contrast to human and animal studies, plant data are often generated from experimental designs in which the experimental units are field plots composed of multiple plants from a common family or inbred line, and often the designs are replicated across different environments. A typical linear model that accounts for environment, genotype, and genotype-by-environment interactions requires multiple random terms, each associated with a different variance component. Although a full model incorporating these random effects in addition to the effect of a single marker can be specified and fit using a mixed linear model, this approach is too computationally demanding for practical use in scanning thousands or millions of markers in a GWAS.

Software such as EMMA (Kang *et al.* 2008), FaSTLMM (Lippert *et al.* 2011), and GEMMA (Zhou and Stephens 2012) were developed to solve the large computational problem in human datasets. EMMA takes advantage of the specific nature of the optimization problem in applying mixed models for association mapping by leveraging spectral decomposition of the genomic relationship matrix. By substantially decreasing the computational cost of each iteration, it enables convergence to a global optimum of the likelihood in variance-component estimation with high confidence by combining grid search and the Newton-Raphson algorithm. Since repeatedly estimating variance components for each SNP is computationally expensive, approximate algorithms like ‘EMMA expedited’ (called EMMAX) and ‘population parameters previously determined’ (called P3D) provide additional computational savings by assuming that variance parameters for each tested SNP are the same (Kang *et al.* 2010; Zhang *et al.* 2010). More recently, FaST-LMM and GEMMA algorithms were proposed that can perform rapid GWAS analysis without assuming variance parameters to be the same across SNPs. FaST-LMM uses spectral decomposition of the genetic similarity matrix to transform (rotate) the phenotypes, SNPs and covariates. These transformed data are uncorrelated can be analyzed with a linear regression model. Similarly, GEMMA expedites each iteration by optimizing the efficiency of the computations required to evaluate the model likelihood and the first and second derivatives of the likelihood function. However, these software provided solutions for linear mixed models that only involve two random components: the polygenic background and error variance components.

In many plant studies, a full accounting of extraneous variation requires multiple random terms, each with a separate variance component, such that two-stage analyses are still necessary even with these improvements in algorithms to conduct linear mixed model GWAS. Two-stage approaches to GWAS for plant studies replicated across environments can take various forms. For example, in the first stage, the genotype effects can be fit as fixed or as random effects with no covariances, leading to the marginal prediction of genotype effects as either best linear unbiased estimation (BLUE) or best linear unbiased prediction (BLUP), respectively. In the second stage, the BLUEs or BLUPs of genotype obtained in the first stage may be fit as the dependent variable in a GWAS, in which the genotypes are treated as random with a variance-covariance matrix proportional to an estimated realized genomic relationship matrix (Aranzana *et al.* 2005; Zhang *et al.* 2009; Peiffer *et al.* 2013; Lipka *et al.* 2013; Pasam *et al.* 2012).

Another approach to two-step GWAS involves using residuals, similar to the GRAMMAR method, but a complication is that replicated trials result in multiple residual values for each family. The first stage residuals could be averaged for each family and used as inputs to the second stage. Alternatively, a term for independent family effects can be fit in the first stage model in addition to the polygenic family effects with covariance proportional to the relationship matrix (Oakey *et al.* 2007) and the independent line effect could be used as the dependent variable in the second stage. Finally, a three-step analysis procedure could be used. In the first step, BLUEs are computed for each line from plot level data, second BLUEs are fit as dependent variables in a linear mixed model including the relatedness matrix, and in the third step, residuals from second step are used for GWAS.

In addition to more complex experimental designs, another common feature of plant datasets is their unbalanced nature. Balanced data sets contain an equal number of observations for each combination of model factor levels. In contrast, plant breeding data sets often involve a series of trials over locations and years in which the genetic entries differ across environments. In addition, some data are often missing due to practical problems, and even within environments, experimental designs are often not balanced. The lack of balance impacts two-stage analyses in several ways. First, the BLUPs of lines that are represented by fewer records in the data set are shrunk back to the population mean to a greater extent than lines with more records. Second, the BLUEs or BLUPs obtained from the mixed model analysis of an unbalanced data set have variable standard errors. The variation in precision among the BLUEs or BLUPs is ignored in the second stage analysis, resulting in a loss of information. Simulation studies (Wang *et al.* 2011) indicate that unbalanced data in two-stage GWAS can cause more false positives.

Methods for analyzing a series of unbalanced performance evaluations of crops have been considered in detail in the context of maximizing the precision and accuracy of marginal predictions of the genetic entries (Smith *et al.* 2009; Möhring and Piepho 2009). In this context, single-step analysis is considered optimal, but may have high computational demand. Two-stage analysis of crop performance trials involves analyzing individual trials separately, then using family BLUEs from each trial as dependent variables in a simplified second stage analysis. Two stage analysis methods that use weighted analysis in the second step, in which weights are proportional to the precision of the BLUEs from the first step, often provide close approximation to the results of a single stage analysis (Smith *et al.* 2009; Möhring and Piepho 2009). Additional complexity in the two stage analysis occurs when the residual values within environments are not independent, as occurs when spatial correlations are modeled in the residual variance structure. This results in lack of independence among the BLUEs; however approximate and exact methods have been developed to account for this lack of dependence as well as the variable precision among BLUEs in the second stage of analysis (Möhring and Piepho 2009; Piepho *et al.* 2012).

Inspired by previous work on two stage analysis of crop performance trials, George and Cavanagh (2015) proposed a two-stage GWAS approach that weights the BLUEs for families from the first stage in the linear mixed model GWAS scan. Their results indicate that the weighted two-stage GWAS provided comparable results to the single stage GWAS, and suggest that weighted two-stage analysis appears is a useful approach for conducting GWAS using data from multi-environment plant breeding trials. Several questions about the use of two-stage GWAS remain unanswered, however. First, it is unclear which summary variable is appropriate to use as a dependent variable for second-stage GWAS (Pasam *et al.* 2012). In particular, the use of BLUP in two-stage analyses in which the hypothesis test is conducted only in the second stage has been criticized (Hadfield et al. 2010).An alternate approach of using residuals from a first stage mixed model accounting for genomic relationships as dependent variables in the second stage may also be considered. Second, to our knowledge, none of the specialized open-source GWAS software packages have the flexibility to incorporate weights in the residual variance structure.

The objective of this study was to compare one-stage and several different two-stage GWAS methods using simulated real marker data and simulated phenotype data from a large maize diversity panel. Different levels of imbalance were imposed on the data to evaluate the effect of data imbalance. False discovery rate and power of marker-trait association tests and estimates of marker effects were compared for six different methods: one-stage analysis, two-stage unweighted analysis based on BLUPs or BLUEs from the first stage, two-stage weighted analysis based on BLUPs or BLUEs, and analysis of residuals after estimating random family effects with the relationship matrix. A weighted two-stage method that incorporates information on the variance of first-stage marginal predictions was implemented in the publicly available software TASSEL (Bradbury *et al.* 2007).

## Material and Methods

### Simulation data set

To reflect the real linkage disequilibrium (LD) structure of genome, the genotype used for simulation is from a subset containing 2480 lines representing almost all of the available inbred maize lines from the USDA Plant Introduction collection (Romay *et al.* 2013). After the initial imputation described in Romay et al. (2013), ~16% of line-marker combinations were still missing. An additional imputation was performed using Beagle 4.0 (Browning and Browning 2009). A subset of 111,282 SNP markers was obtained by filtering out markers that have estimated imputation accuracy less than 0.995 and pairwise genotypic correlation greater than 0.5 by linkage-disequilibrium pruning using PLINK (Purcell *et al.* 2007). Data for *g* = 2480 inbred lines and *n_env_* = 10 environments was simulated.

For each simulation data set, *q* = 10 or *q* = 50 SNPs were randomly sampled from among markers with minor allele frequency greater than or equal to 0.01, with the restriction that no pairs of markers were within 20 adjacent marker positions to avoid high LD between QTL. Markers selected as causal loci were assigned a constant QTL effect, other markers had zero effect. Genotypic values were created by simulating both QTL and polygenic background effects. The phenotype was simulated as the sum of major gene effects, polygenic genetic background, environmental effect and random error, for *i* = 1 to *g* lines, *j* = 1 to *n_env_* environments and *k* = 1 to *q* QTL (*q* = 10 or 50).:

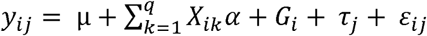

where polygenic effect 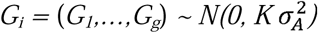, environmental effect 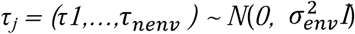, and random error 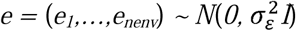, *X*_ik_ are the coefficients for QTL effects, reflecting the number of copies (0, 1, or 2) of the minor allele at each QTL in line *i*, and α is the effect of each QTL (which we set constant for all QTL within one replication). By changing 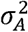,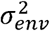,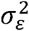 and α we were able to simulate a reasonable range of heritabilities (Table 1).

**Table 1.** Parameter settings for simulation study.

| Experimental design | QTL number | QTL effect | $\sigma_A^2$ | $\sigma_{env}^2$ | $\sigma_\varepsilon^2$ | Proportion of genotypic variance due to all QTL | Average proportion of genotypic variance due to one QTL (%) | Total heritability | Heritability associated with QTL effects | Heritability associated with polygenic effects | Heritability accounted by each QTL (%) |
| --- | --- | --- | --- | --- | --- | --- | --- | --- | --- | --- | --- |
| Balanced | 10 | 11 | 109 | 100 | 2500 | 0.83 | 8.3 | 0.85 | 0.69 | 0.47 | 7 |
| Balanced | 10 | 6.6 | 289 | 100 | 2500 | 0.48 | 4.8 | 0.85 | 0.41 | 0.74 | 4 |
| Balanced | 50 | 5.2 | 105 | 100 | 2500 | 0.86 | 1.7 | 0.86 | 0.74 | 0.46 | 1.5 |
| Random unbalanced | 10 | 11 | 109 | 100 | 2500 | 0.83 | 8.3 | 0.74 | 0.61 | 0.39 | 6 |
| Random unbalanced | 10 | 6.6 | 289 | 100 | 2500 | 0.48 | 4.8 | 0.74 | 0.36 | 0.61 | 3.6 |
| Random unbalanced | 50 | 5.2 | 105 | 100 | 2500 | 0.86 | 1.7 | 0.75 | 0.65 | 0.40 | 1.2 |
| Severely<br>unbalanced | 10 | 11 | 109 | 100 | 2500 | 0.83 | 8.3 | 0.58 | 0.49 | 0.30 | 5 |
| Severely<br>unbalanced | 10 | 6.6 | 289 | 100 | 2500 | 0.48 | 4.8 | 0.59 | 0.30 | 0.50 | 3 |
| Severely<br>unbalanced | 50 | 5.2 | 105 | 100 | 2500 | 0.86 | 1.7 | 0.60 | 0.51 | 0.30 | 1 |

We simulated three different genetic architectures varying for the number of QTL and QTL effect sizes: 10 QTL accounting for 83% of total genotypic variation, 10 QTL accounting for 43% of genetic variation, and 50 QTL accounting for 86% of genotypic variation (Table 1). The proportion of variance associated with QTL was estimated as the correlation between the sum of QTL effects and the phenotypic value of each line. The proportion of variance associated with polygenic background effects was estimated as the correlation between the polygenic effects and the phenotypic effects. The QTL and polygenic effects were not independent, so the total heritability was generally less than the sum of the QTL and polygenic variances. Furthermore, we assigned constant effects to all QTL within a genetic architecture setting, but since the QTL were randomly sampled from the true markers, their allele frequencies varied and the heritability due to QTL also varied among datasets. Hereafter, we refer to the average proportion of total heritability explained by QTL across datasets when this proportion is indicated, which means that heritability associate with each QTL is calculated as the average heritability accounted by each QTL.

We simulated three different scenarios for missing data: complete balanced data, randomly missing unbalanced data, and severely unbalanced data (Table 1). Balanced datasets had all lines evaluated at all environments with no missing values (24800 records). The two unbalanced datasets were generated from each complete dataset. Randomly unbalanced datasets contained a random subset of 50% of the data of the complete dataset (12400 records). Severely unbalanced datasets had half of the lines evaluated at only one environment and the other half of lines evaluated at ten environments (13640 records). We generated 50 replicate complete data sets for each genetic architecture and two random subsets of each complete data set (100 replicates total) for each unbalanced data setting.

The realized additive genomic relationship matrix was estimated using R software version 3.0.0 (R Core Team 2013) based on observed allele frequencies (VanRaden 2008; method 1). The dataset for calculating relationship matrix is the whole genotype dataset.

### Analysis methods

The simulated datasets were analyzed using each of six methods (Table 2).

**Table 2.** GWAS methods used.

| Method | First stage<br>Model | First stage<br>distribution of $F_j$ | Summary result of first<br>stage | Second stage model | Second stage<br>distribution of $F_j$ | Summary result of<br>second stage |
| --- | --- | --- | --- | --- | --- | --- |
| One-stage | $Y_{ijk}$<br>$= \mu + E_i + F_j$<br>$+ X_{jk}\beta_k + \varepsilon_{ijk}$ | $F_j \sim N(0, K\sigma_A^2)$ | Effect of marker $k = \hat{\beta}_k$ | | | |
| Unweighted<br>BLUE 2-stage | $Y_{ij}$<br>$= \mu + E_i + F_j$<br>$+ \varepsilon_{ij}$ | Fixed effect | $BLUE(Y_{.j}) = \mu + \hat{F}_j$ | $BLUE(Y_{.j})$<br>$= \mu + F_j + X_{jk}\beta_k$<br>$+ \varepsilon_{jk}$ | $F_j \sim N(0, K\sigma_A^2)$ | Effect of marker $k = \hat{\beta}_k$ |
| Unweighted<br>BLUP 2-stage | $Y_{ij}$<br>$= \mu + E_i + F_j$<br>$+ \varepsilon_{ij}$ | $F_j \sim N(0, I\sigma_f^2)$ | $BLUP(Y_{.j}) = \mu + \hat{F}_j$ | $BLUP(Y_{.j})$<br>$= \mu + F_j + X_{jk}\beta_k$<br>$+ \varepsilon_{jk}$ | $F_j \sim N(0, K\sigma_A^2)$ | Effect of marker $k = \hat{\beta}_k$ |
| Weighted<br>BLUE 2-stage | $Y_{ij}$<br>$= \mu + E_i + F_j$<br>$+ \varepsilon_{ij}$ | Fixed effect | $BLUE(Y_j) = \mu + \hat{F}_j$<br>$w_j = V(\hat{Y}_j)$ | $BLUE(Y_j)$<br>$= \mu + F_j + X_{jk}\beta_k$<br>$+ \varepsilon_{jk}$<br>$\varepsilon_{jk} \sim N(0, \mathbf{I}w\sigma_\varepsilon^2)$ | $F_j \sim N(0, \mathbf{K}\sigma_A^2)$ | Effect of marker $k = \hat{\beta}_k$ |
| Weighted<br>BLUP 2-stage | $Y_{ij}$<br>$= \mu + E_i + F_j$<br>$+ \varepsilon_{ij}$ | $F_j \sim N(0, \mathbf{I}\sigma_f^2)$ | $BLUP(Y_j) = \mu + \hat{F}_j$<br>$w_j = V(\hat{Y}_j)$ | $BLUP(Y_j)$<br>$= \mu + F_j + X_{jk}\beta_k$<br>$+ \varepsilon_{jk}$<br>$\varepsilon_{jk} \sim N(0, \mathbf{I}w\sigma_\varepsilon^2)$ | $F_j \sim N(0, \mathbf{K}\sigma_A^2)$ | Effect of marker $k = \hat{\beta}_k$ |
| Residual 3-stage | Step 1:<br>$Y_{ij} = \mu + E_i + F_j + \varepsilon_{ij}$<br>Step 2:<br>$BLUE(Y_j)$<br>$= \mu + F_j + \varepsilon_j$ | Step 1: Fixed effect<br>Step 2:<br>$F_j \sim N(0, \mathbf{K}\sigma_A^2)$ | Step 1: $BLUE(Y_j) = \mu + \hat{F}_j$<br>Step 2:<br>$resid(Y_j) = \varepsilon_j$ | $resid(Y_j)$<br>$= \mu + X_{jk}\beta_k + \varepsilon_{jk}$ | NA | Effect of marker $k = \hat{\beta}_k$ |

### One-stage model analysis

Suppose that *n* total observations were made on *g* lines so that ***Y*** is an *n* × *1* vector of observed phenotypes. A linear mixed model for single-stage association mapping is expressed as:

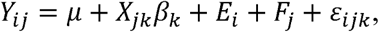

where *Y_ij_* is the observation on family *j* in environment *i*, *E_i_* are random macro-environment main effects, *X_jk_* is the genotype score (0, 1, or 2 indicating number of minor alleles) of family *j* at marker *k*, *β_k_* is the fixed effect of marker *k*, *F_j_* are random genetic background effects, and ε*_iJk_* are residual effects. The distributions of random effects are 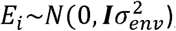,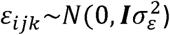 and 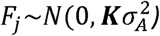, where ***K*** is the *g* × *g* kinship matrix inferred from genotypes based on observed allele frequencies ((VanRaden 2008), methodl). Since estimating the variance components, particularly 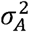, is computationally intensive, we used the parameters previously determined method for the GWAS scan. For each simulation data set, the variance components were estimated once by restricted maximum likelihood from a reduced model with no fixed marker effects using ASReml. The variance components were then fixed at those values while subsequently testing each marker (Zhang *et al.* 2010). After obtaining estimates of 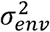,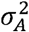,and 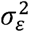, and using ASReml, the effect of marker *k* was estimated as:

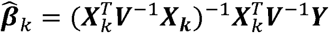

In this formula, **β***_k_* is a vector of the fixed intercept, μ, and one or two genotypic effect estimates for marker *k*. Markers with two genotypic classes require only one genotypic effect estimate, whereas markers with three genotypic classes require two effect estimates. ***X_k_*** is an *n* × 2 or *n* ×3 matrix consisting of a column of ones and one or two columns of dummy variables (depending on the number of genotypes at the marker) indicating the different genotypes at marker *k*. Markers with two genotypic classes require one column, whereas markers with three genotypic classes require two columns of dummy variables in 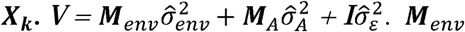, is a block diagonal *n* × *n* matrix, indicating environmental effect correlations of 1 for observations within a common environment and 0 for pairs of observations in different environments. Each block is a *g_i_*×*g_i_* matrix where every element is 1 and *g_i_* is the number of genotypes evaluated in environment *i.* ***M_env_*** can be constructed as 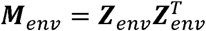, where ***Z****_env_* is the *n* × *e* design matrix for environment effects (*e* = 10 in all of our data sets). **M**_A_ is an *n* × *n* matrix, where each element is the realized genomic relationship coefficient for a pair of observations. **M**_A_ can be constructed as 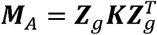, where ***Z****_g_* is the *n* × *g* design matrix for inbred line effects (*g* = 2480 in all of our data sets) and **K** is the realized genomic relationship matrix. Note that this analysis requires inversion of **V** but the inverse can be computed once for a given **Y** vector and used for all marker tests in that data set. F-tests were used to test the null hypotheses of zero effect at each marker separately (Yu *et al.* 2006; Kang *et al.* 2008; Kennedy *et al.* 1992). Additive effects of markers were estimated as the appropriate linear combination of homozygous genotype effects.

### Unweighted two stage analysis- BLUP

In the first stage, best linear unbiased predictions (BLUPs) for lines are calculated, assuming independence among the lines:

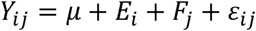

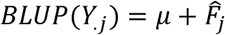

where 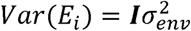,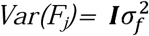,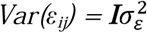.

In the second stage, the *g* × 1 vector of line BLUPs are treated as the dependent variable in the marker tests:

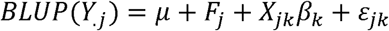

where 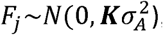, ***X***is a *g* × *q* design matrix for marker effects, *β* is a *q* × 1 vector representing coefficients of the fixed marker effects, and 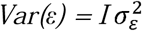. The variance components 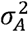 and 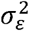 are estimated once without fixed marker effects, and the resulting estimates used as fixed values in subsequent tests of each marker.

### Unweighted two stage analysis- BLUE

In the first stage, best linear unbiased estimates (BLUEs) of lines are calculated:

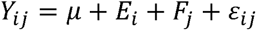

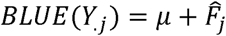

where 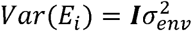,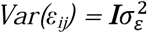, and *F_j_* is treated as a fixed effect

In the second stage, the *g* × 1 vector of line BLUEs is treated as the dependent variable:

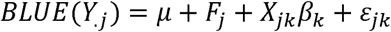

where 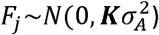, *X* is an *g* × *q* design matrix for marker effects, *β* is a *q* × *1* vector representing coefficients of the fixed marker effects, and 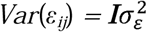. The variance components 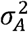 and 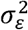 are estimated once without fixed marker effects, and the resulting estimates used as fixed values in subsequent tests of each marker.

### Weighted –two stage analysis (BLUE and BLUP)

BLUEs or BLUPs are calculated in the first step, and the variance of each BLUE or BLUP is also recorded for use in the second step. The second step fits the following model:

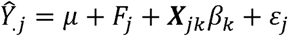

where *Ŷ_.j_* is a *g* × 1 vector of BLUE or BLUP values for *g* lines, 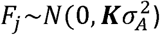, **X** is a *g* × *q* design matrix for marker effects, *β* is a *q* × *1* vector representing coefficients of the fixed marker effects, and the distribution of residual effects is: 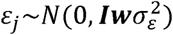,***w****_j_* = *V*(*Ŷ_.j_*). Thus, the weighted two-stage analyses differ by weighting the diagonal elements of the residual variance-covariance matrix with the variances of the BLUEs or BLUPs. The variance components 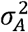 and 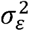 are estimated once without fixed marker effects, and the resulting estimates used as fixed values in subsequent tests of each marker.

### Residual three stage

BLUEs are calculated as described above, then the following model is fitted in the second step:

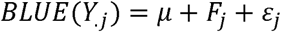

where 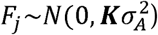 and 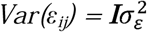. Then the vector of residuals, ε, is used as the dependent variable in the last step analysis:

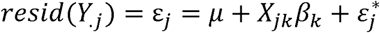

where ***X***is a *g* × *q* design matrix for marker effects, *β* is a *q* × *1* vector representing coefficients of the fixed marker effects, 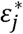 are residuals from the final stage model and are distinguished from the line residuals from the second stage, and 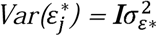. Each marker was tested separately in the final stage.

In multiple-step models, the initial stage linear mixed models were fitted used ASReml 3.0 software. Single-step analysis marker scans were conducted using a custom Python script that uses the variance components for the current data set estimated without fixed marker effects with ASReml, computes **V**^-1^, and uses that **V**^-1^ for testing each marker. The marker scan steps for other analyses were conducted using TASSEL (Bradbury *et al.* 2007). We implemented the weighted two-stage marker scan as an option in TASSEL.

### Power and false discovery rate

Significant association tests were declared based on an empirical false discovery rate estimated for each analysis separately using the “qvalue” package in R (Bass *et al.* 2015). Markers with *q*-values less than or equal to 0.05 were treated as significantly associated with the simulated traits. Power was calculated as the ratio of number of true positive association tests to the total number of true QTLs. We then computed the true false discovery rate for each analysis as the proportion of false positive discoveries among all positive discoveries. False positives were defined as significant markers with small linkage disequilibrium (LD) *r*^2^ values with true QTL. We evaluated two different LD thresholds to declare false positives: *r*^2^ < 0.1 and *r*^2^ < 0.05. At each of these thresholds, false discovery rate was calculated as the number of false positive association tests divided by the total number of positive (significant) tests.

### Bias and mean square error of QTL effect estimation

Bias (*b_c_*) and mean square error (MSE) of effect estimates at causal loci were calculated as follows:

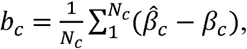

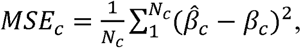

where *N_c_* is the number of causal loci, 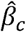 is the estimated additive marker effect for a causal locus and *β_c_* is the true additive marker effect of causal locus.

## Results

### Power and false discovery rate

We simulated three different genetic architectures based on the observed allele frequencies, population structure, and linkage disequilibrium of a large panel of diverse maize inbred lines (Romay *et al.* 2013). The genetic architectures differed by the number and effect size of QTL (Table 1). Within one simulation data set, the QTL effects were constant, but the variation caused by each QTL differed because the allele frequencies differed among the randomly sampled SNPs chosen to represent QTL. The total genetic variation caused by QTL was nearly constant across replicated data sets for a given genetic architecture, however. Therefore, we characterized the genetic architectures by the average genetic variance associated with one QTL, which varied from 1.7% for the situation of 50 QTL to 8.3% for the 10 QTL with large effects (Table 1). The average total heritability (due to both QTL and background effects) for all three genetic architectures was consistent, varying only between 85% and 86% (Table 1). The sum of heritability due to QTL and heritability due to polygenic effects was larger than total heritability. This occurred because the QTL are not independent of the genetic background effects. As a result, the variance due to line polygenic background effects cannot be entirely separated from the variance due to QTL. The correlation between polygenic effects and total genetic value is influenced by the correlated effects of QTL, and vice-versa. This reflects a realistic and characteristic aspect of genetic architecture in populations with substantial structure and local LD.

For each complete simulated data set, we generated four additional subsets, reflecting two replicated samplings of two different missing data patterns. In addition to the variability for QTL effects across replicated simulation data sets for a common genetic architecture, the different missing data patterns resulted in variability in the total heritability of line means. On average, however, the total heritabilities were very similar across genetic architectures for a given missing data pattern. On average,50% randomly missing data resulted in a line mean heritability of about 75%, and the severely unbalanced situation resulted in a line mean heritability of about 60% (Table 1).

For balanced datasets, power of association tests was similar for two-stage analyses using BLUEs and BLUPs in the second stage (Figure 1). When data are balanced, the variance of BLUEs and BLUPs from the first stage of a two-stage analysis are homogeneous, weighted and unweighted second stage analyses are identical. For randomly unbalanced datasets, weighted and unweighted two-stage analysis using either BLUEs or BLUPs were not identical, but had similar power to detect associations (Figure 1). The largest differences among power of different analyses was observed with severely unbalanced datasets, where the weighted BLUE two-stage method had power about equal to the one-stage analysis, weighted and unweighted two-stage analyses using BLUPs were almost as good, but the unweighted two-stage analysis using BLUEs had a notable reduction in power. Analysis using the residuals from a model fitting genetic relationships in a previous step had lowest power among all analyses in all three experimental designs (Figure 1, Table S1). Power of association tests had strong non-linear relationships with minor allele frequency, QTL effect size, and the proportion of missing data (Figure S1).

**Figure 1.**
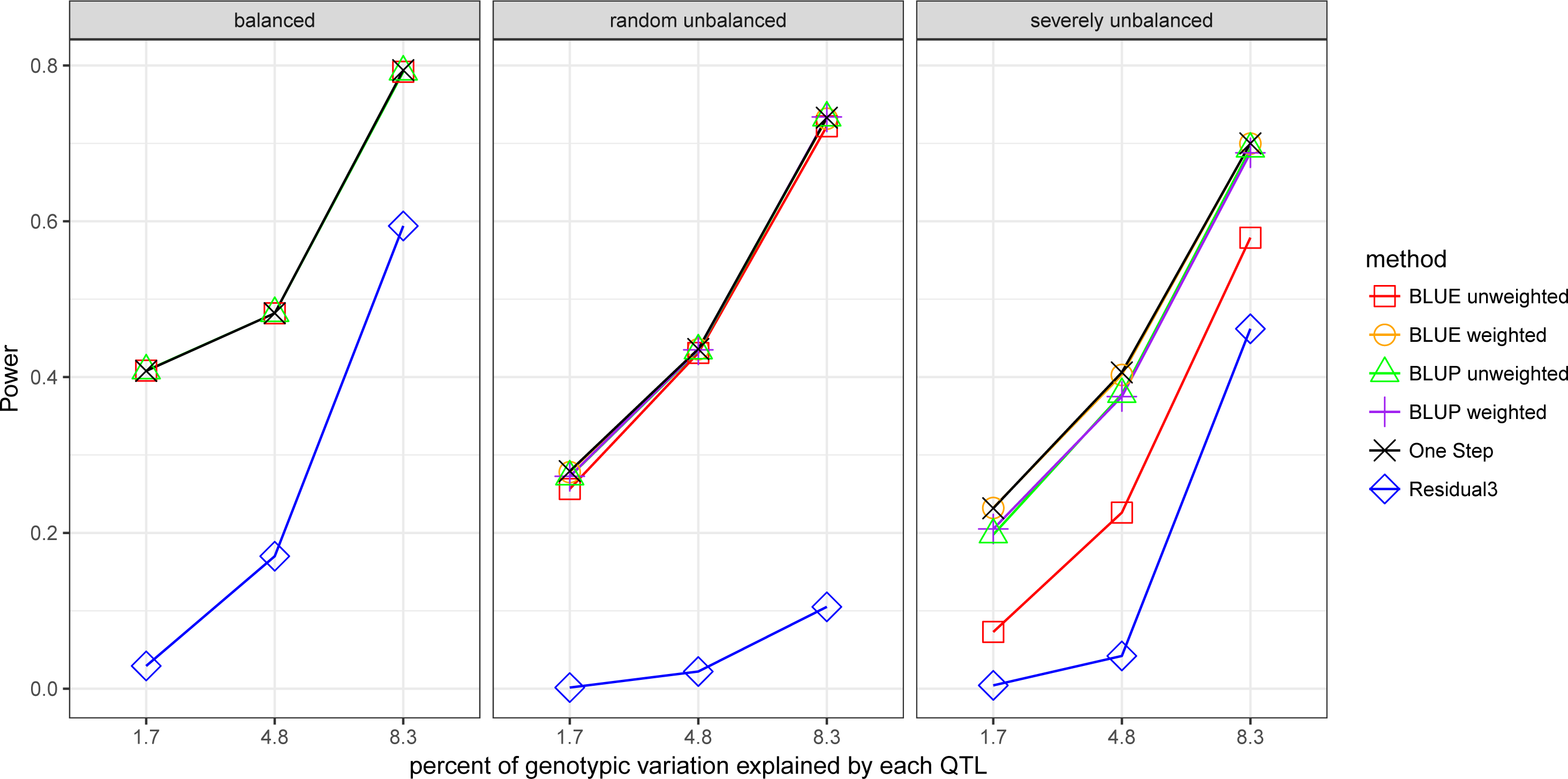
Power of six different association testing methods to detect causal variants for three different genetic architectures and three levels of data imbalance. Balanced datasets had all lines evaluated at all environments with no missing values (24800 records). Randomly unbalanced datasets contained a random subset of 50% of the data of the complete dataset (12400 records). Severely unbalanced datasets had half of the lines evaluated at only one environment and the other half of lines evaluated at ten environments (13640 records).

False discovery rate (FDR) was similar for one-stage and all two stage analyses (using BLUEs or BLUPs and weighted or not; Figure 2, Table S2). The three-stage analysis based on residuals had very poor power and resulted in very few positive discoveries (Table S1), so this method is not included in the comparisons of false discovery rate. All methods had an inflated FDR (actual FDR greater than the estimated rate) for the simplest genetic architecture (10 QTL accounting for 83% of true genetic variance; Figure 2). The FDR inflation was most severe for balanced data (FDR about 15%), but still exists for the unbalanced cases (Figure 2). We excluded markers with LD *r*^2^ > 0.1 with causal QTL for the analysis reported in Figure 2. When we restricted the computation of FDR to only markers LD *r*^2^ < 0.05 with QTL, FDR for balanced data and simplest genetic architecture dropped to about 7% (Figure 3). The strong dependency of FDR on LD with QTL in in this case indicates that even low levels of LD with causal markers can inflate FDR when the QTL effects are large.

**Figure 2.**
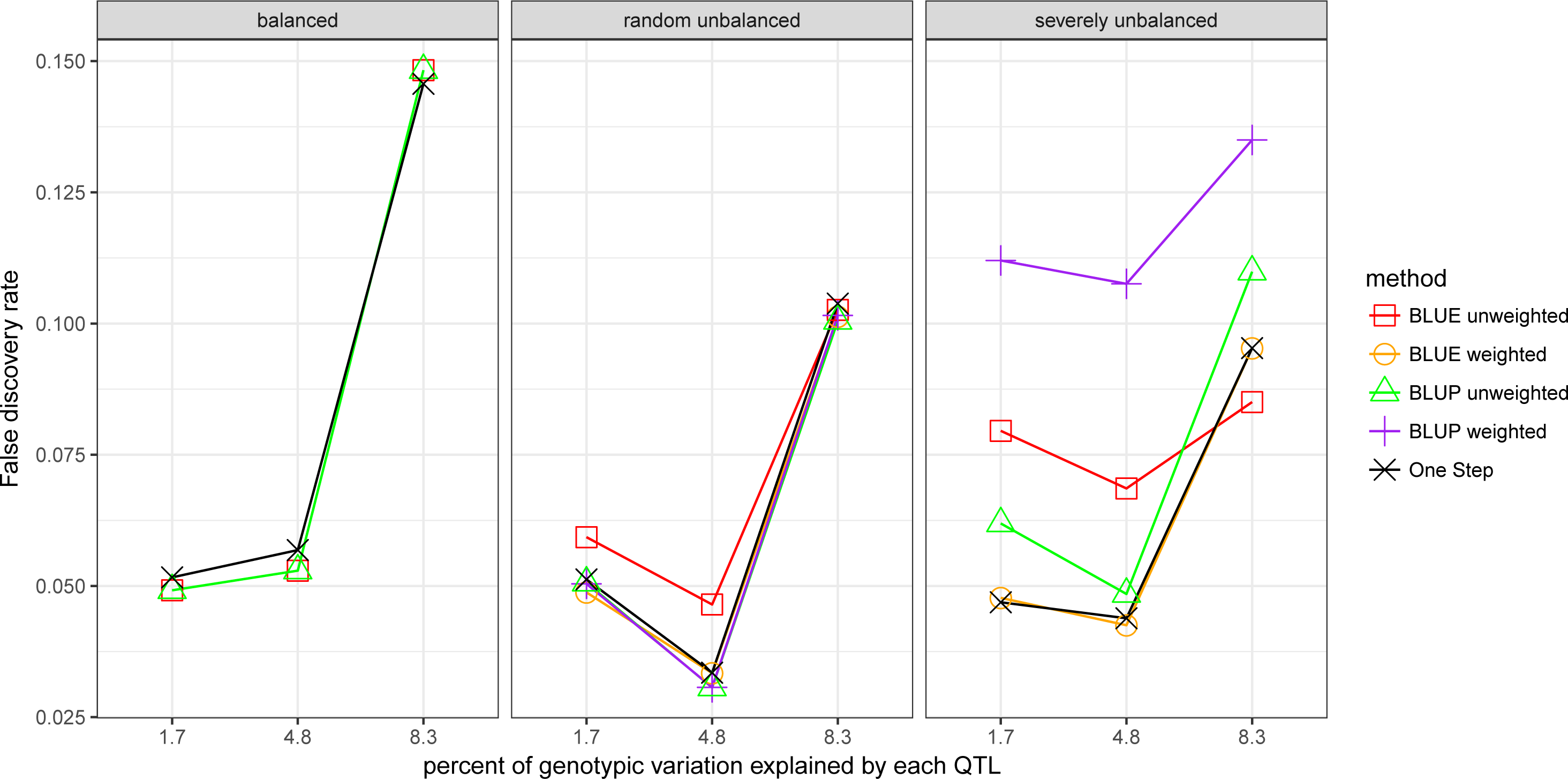
False discovery rate when false positives were defined as markers that had LD *r*^2^ < 0.1 with true QTL and were declared significant at the empirically estimated *q* < 0.05. False discovery rate is the proportion of false positives defined this way among all markers declared significant. False discovery rate for residual 3-step method is not shown, since it identified very few significant markers.

**Figure 3.**
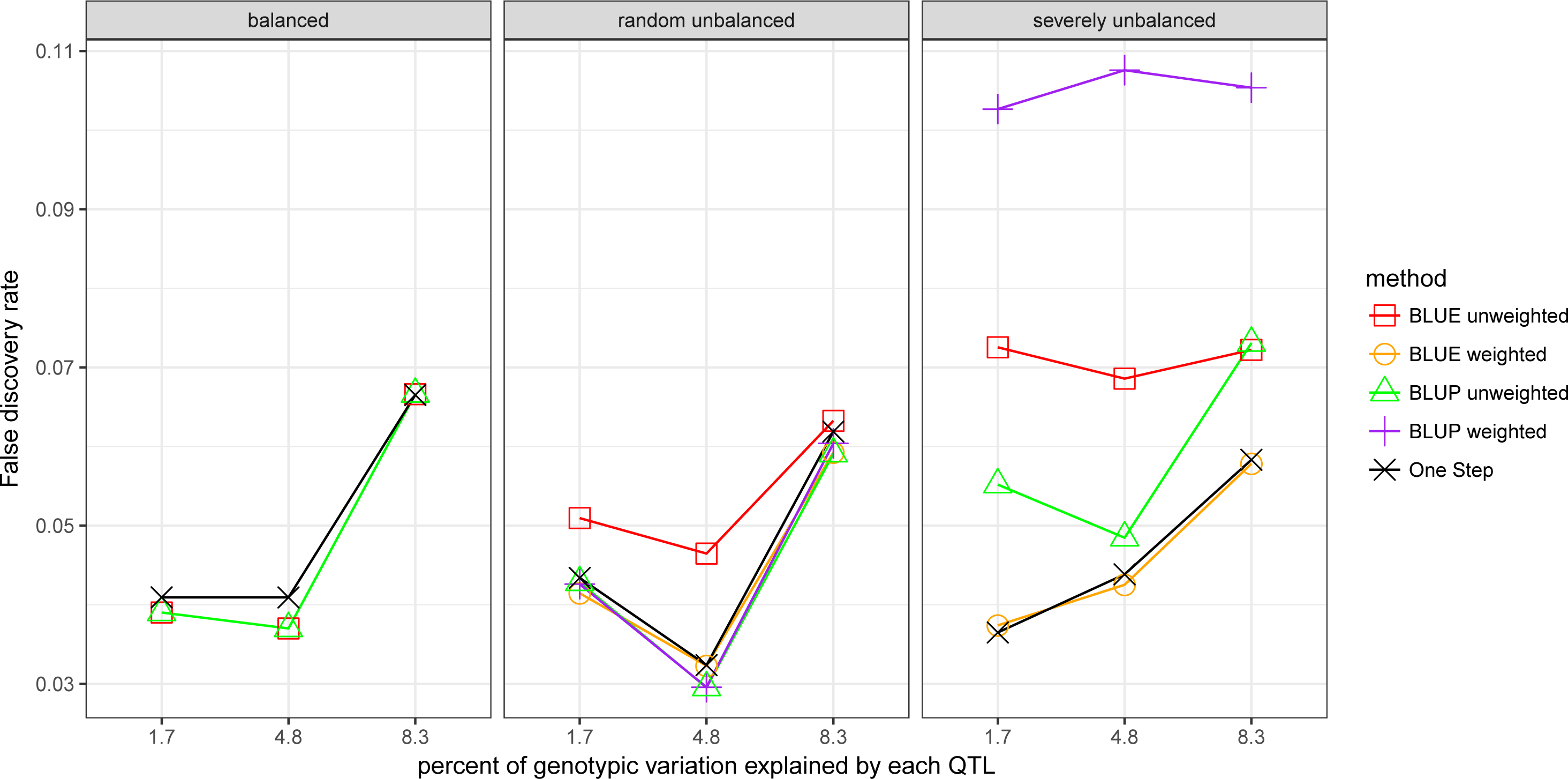
False discovery rate when false positives were defined as markers that had LD *r*^2^ < 0.05 with true QTL and were declared significant at the empirically estimated *q* < 0.05. False discovery rate is the proportion of false positives defined this way among all markers declared significant. False discovery rate for residual 3-step method is not shown, since it identified very few significant markers.

The one-stage and weighted BLUE two-stage analyses had the best FDR when data were severely unbalanced (Figures 2 and 3). The unweighted BLUE two-stage analysis had higher FDR than the weighted BLUE two-stage analysis in most unbalanced conditions. Weighted BLUP two-stage analysis had similar FDR to one-stage for balanced and randomly unbalanced data, but had dramatically inflated FDR with severely unbalanced data. The weighted BLUP two-stage method had worse FDR than all other methods, including the unweighted BLUP method, when data were severely unbalanced. This effect remains even when FDR was computed for only markers with LD *r*^2^ < 0.05 with QTL (Figure 3), and the inflation is strong even in the most polygenic architecture, so it is not simply a function of LD with causal QTL.

Because the power and FDR results suggest that the weighted (but not unweighted) BLUE two-stage analysis has similar properties to the one-stage analysis, we compared the distribution of genome-wide association test *p*-values for the weighted and unweighted BLUE two-stage methods to the one-stage analysis. The distribution of *p*-values for all SNP tests was nearly identical for the one-stage and weighted BLUE two-stage methods (Figure 4). The severely unbalanced data case in particular results in many large deviations of unweighted BLUE two-stage *p*-values compared to the one-stage *p*-values, however (Figure 4).

**Figure 4.**
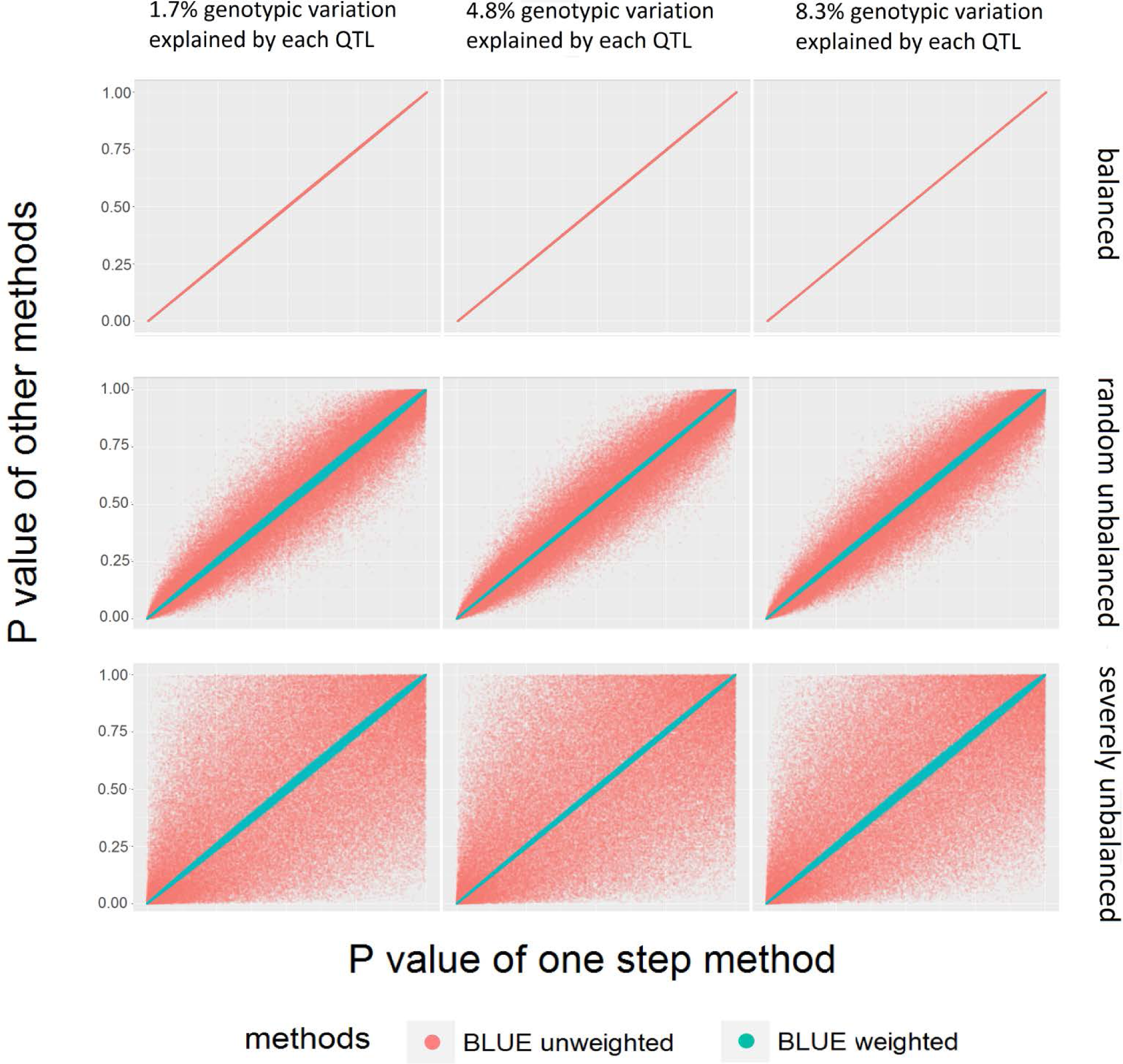
Distributions of all genome-wide marker association test *p*-values using the one-step analysis (Y-axis) or two-step analysis (X-axis) for three different genetic architectures and three levels of data imbalance. *P*-values from weighted BLUE two-stage method are in blue and *p*-values from unweighted BLUE two-stage method are in orange.

**Figure 5.**
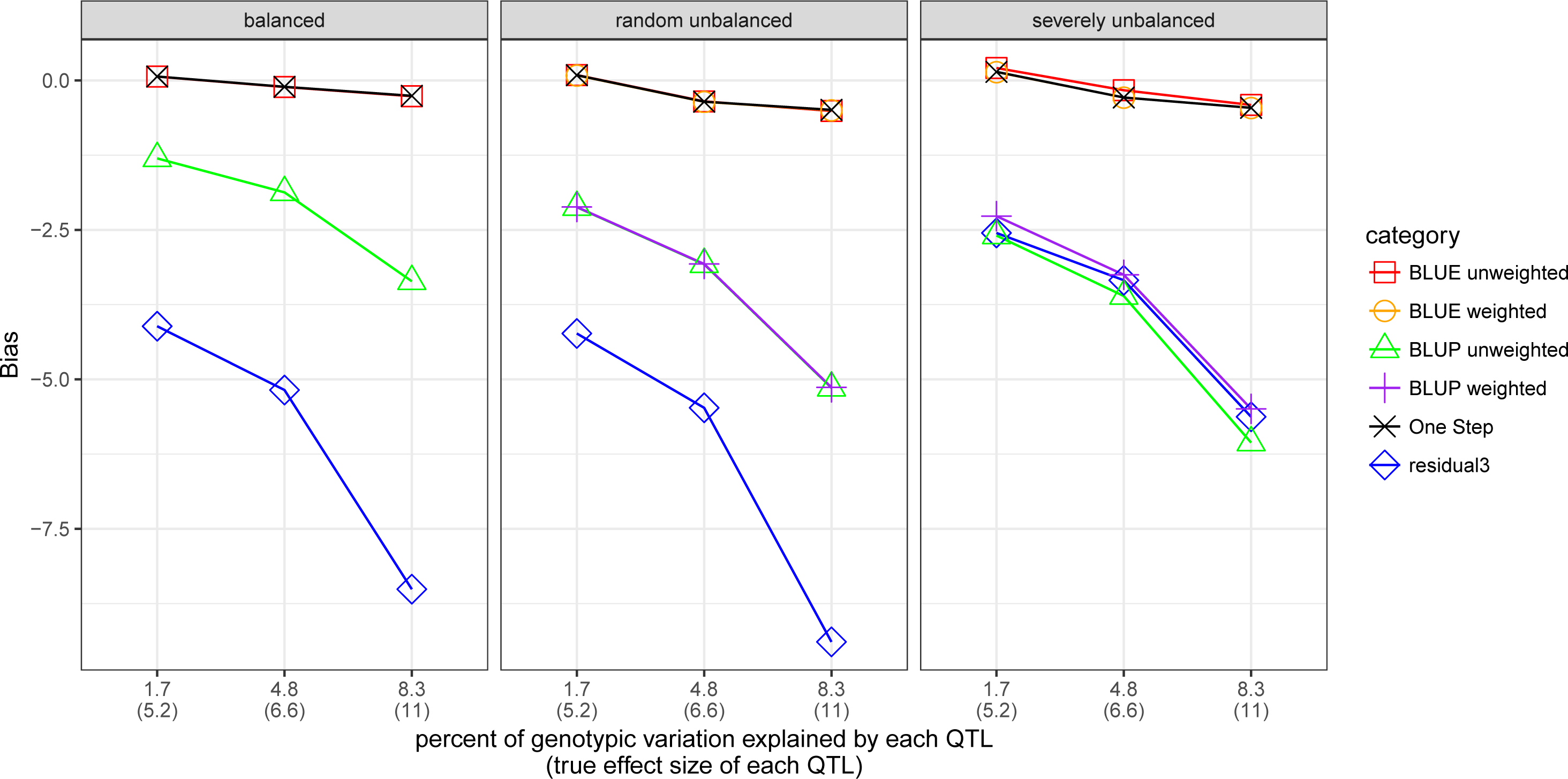
Bias of QTL effect estimates from different GWAS methods for three different genetic architectures and three levels of data imbalance.

### Bias and MSE

Estimated effects of true QTL are biased downward for all methods except the one-stage and the two-stage BLUE methods (Figure 4). The residual 3-stage method results in the strongest downward bias under most combinations of genetic architecture and data structure. Methods using BLUPs also have downward bias under all conditions, as a result of the shrinkage of line values that occurs before the final step GWAS scan. For example, for the severely unbalanced data case and large-effect QTL effect, the weighted and unweighted BLUE two-stage analyses estimated the QTL effects with a bias of −0.5 units,or −4% of the true value, whereas the two-stage BLUP methods had downward bias around −6 units, around 50 % of the true value. These trends are also reflected in the mean square error variance for effect estimates (Figure S2; Table S3).

### Computational time

For a single analysis analyzing 2480 lines with ~110,000 markers using a single core (Intel Xeon E5-2680v3), one-step analysis required 146 hours, whereas the GWAS scan (second step) of two-stage methods required 30 hours for unweighted and 32 hours for weighted methods. The two-stage analyses also involved a first step to estimate BLUEs or BLUPs, which required 0.3 hours. Three-stage analysis using residuals required around 10 hours for the GWAS scan, in addition to 0.4 hours for the first two steps of the analysis.

## Discussion

Our simulation used the real marker data on a large and diverse maize inbred line panel with substantial population structure. For each simulation data set, we assigned a very small proportion of markers to have true causal effects, allowing us to test power of association tests directly at causal variants. In real GWAS studies, however, the researcher cannot assume that the causal variants have been genotyped and included in the marker data set. Instead, researchers rely on sufficient marker density and linkage disequilibrium to detect association signals at markers physically linked in close proximity to causal variants, while trying to reduce the influence of longer-range (and unlinked) LD due to population structure on association tests. In general, linkage disequilibrium decays rapidly in diverse maize panels. On average, LD is below *r*^2^ = 0.2 for markers separated by more than one kb, but there is a large variance around this average value, such that a small proportion of distantly separated marker pairs may still have high LD (Romay *et al.2013*)

Therefore, the power of association tests reported in Figure 1 and Table S1 represents the optimal but unrealistic situation of having the casual variants in the marker data set. Estimating power at markers linked to causal variants introduces some complication into the concept of power, because different researchers have different criteria for considering an association to be a true positive result or a false positive result, depending on how close the marker is to the true variant in physical or genetic distance. Power of association tests reported here was somewhat lower than power for detecting QTL accounting for similar proportions estimated from a simulation study of the maize nested association mapping (NAM) design (Yu *et al.* 2006). The lower power of QTL detection in a diversity panel than in a balanced multiple biparental family design like NAM is expected, as the NAM panel has a simple, known population structure that can be accounted for in the analysis, and more balanced allele frequencies. Power in this study was strongly related to allele frequency, power increased sharply in most cases for minor allele frequencies between 0.05 and 0.20 (Figure S1). Below 5% minor allele frequency, power was very low (~ 0.12) except for the largest-effect QTL simulated (Figure S1). Missing data reduced power as expected, and the effect was greater for severely unbalanced data, even though the total proportion of missing data in that case is not highest (Figure S1). The power of association tests reported here also reflect a rather large sample of inbred lines (2480), which is larger than many association panels currently studied. Thus, in smaller panels, power of detection will be lower than that reported here.

The simulation results clearly demonstrate that the ‘GRAMMAR’ method for conducting a GWAS scan on residuals from a model including random family effects with covariances proportional to the estimated realized genomic relationship coefficients has worse performance (lower power and higher bias) than other methods evaluated in this study. A similar result was reported by Zhou and Stephens (2012) based on a comparison between results of GEMMA, EMMAX and GRAMMAR algorithm scans of a human genetics data set. The particularly poor performance of GRAMMAR in our simulation is likely related to numerous close relationships among lines in the collection of maize inbreds studied, coupled with a preponderance of low minor allele frequencies. In this situation, the QTL effects may tend to be restricted to relatively few groups of closely-related lines, and therefore mostly absorbed into the polygenic background effects.

The two-stage methods using line BLUPs from an initial analysis that regards line effects as random but independent had power of association tests about equal to the one-stage analysis (Figure 1). Weighting had almost no effect on the power or bias of two-stage BLUP methods (Figures 2, 3, and 4). However, both BLUP methods had considerable bias in estimation of QTL effects, due to the shrinkage of line values that are used as dependent variables in the GWAS scan. In general, the unweighted BLUP two-stage method performed similar to or better than the weighted two-stage BLUP method, because the weighted BLUP two-stage had considerably inflated FDR when data were severely unbalanced (Figures 2 and 3). This may have occurred because BLUP itself introduces shrinkage toward the mean of the line values, and the shrinkage is greatest for lines with most missing data. Weighting during the GWAS then decreases the relative influence of lines with highest prediction error variance, which are the same lines whose values have shrunk most toward the mean. The double action of shrinking and underweighting the values of lines with least data increases false discoveries. This can happen when, by chance, lines carrying a rare SNP have complete data, whereas much of the rest of the population (half of lines in our simulation) has a large proportion of missing data. Since the SNP alleles are not independent of the background genetic effects, a subgroup carrying the rare allele but having (by chance) little missing data will reflect the average polygenic effect of the subgroup (even though the line relationships were not accounted for in the model). The line BLUPs in such a subgroup are less likely to be shrunk toward the mean, and, in addition, they have higher relative influence on the association test than other lines. The combination of SNP frequencies correlated with polygenic effects and differential shrinkage and weighting between allelic classes may cause SNP association tests to absorb polygenic effects and produce false positive discoveries.

Users should be cautioned against making inferences about heritability from the relative proportion of genetic and residual variances estimated in the second step of a two-step analysis. The weighting changes the scaling of the residual variance component that is estimated. Therefore, the relative magnitude of the genetic and residual variance components is influenced both by heritability and the scale of the weighting factor.

Our simulation assumed no covariances among the BLUEs computed in the first stage of the analysis. In practice, however, BLUEs may be estimated from complex unbalanced designs such as incomplete block designs, or using models involve spatial correlations among the residuals in first step, leading to correlations among the resulting BLUEs. Various approximate weighting methods have been proposed to handle this situation (Smith *et al.* 2009; Möhring and Piepho 2009; Piepho *et al.* 2012). Piepho *et al.* (2012) also developed a method for exact two-stage analyses, and such a method might also be implemented for GWAS studies, as it has for genomic prediction analysis (Schulz-Streeck *et al.* 2013). In many cases, however, a simple weighting method like the one used here is sufficient to recover the properties of a single-stage analysis (Möhring and Piepho 2009).

Our results suggest that the weighted BLUE two-stage analysis can be recommended across a range of genetic architectures and missing data structures. The power, FDR, and bias of weighted BLUE two-stage analysis was very similar to the one-stage analysis, but with substantially reduced computing time. The method has been implemented in version 5 of the publicly available open-source software TASSEL, available from http://www.maizegenetics.net/tassel. Users need to add an additional file containing the variances of the line BLUEs to the usual two-stage analysis work flow. The file containing the variances of the BLUEs has the same format and header as a TASSEL trait file (so, the same as the file containing the BLUEs themselves). Multiple phenotypes are allowed but they must share the same header and have exactly the same genotype (“taxa”) order. In the TASSEL GUI interface, users must first load the data from four files containing BLUEs, variances of BLUEs, genotype scores, and the relationship matrix, respectively. The BLUE and genotype score data need to be joined using “Intersect Join”. Then, selecting the joined phenotype and marker score data set, the kinship data set and the BLUE variances data set together, the user can choose “weighted MLM” in analysis tab to perform the analysis. To run the analysis from the command line, the same four files are required. An example of the use of command line execution of a weighted analysis using TASSEL version 5 is provided in File S1.

## DATA AVAILABILITY

All data and software codes used to generate simulation data sets and conduct analyses are available for download at: https://drive.google.com/a/ncsu.edu/file/d/0B7Iwvphs9t5hdkdvWUUyVUY0a2M/view?usp=sharing

If the paper is accepted for publication, we plan to post these files permanently at Dryad or a similar public database.

## ACKNOWLEDGMENTS

S.X. was supported by National Institutes of Environmental Health Sciences training grant T32 ES007329 to the North Carolina State University Bioinformatics Research Center and National Science Foundation (NSF) award I0S-1127076; J.B.H. was supported by NSF awards I0S-1127076 and I0S-1238014 and by the U.S. Department of Agriculture, Agricultural Research Service. This work used the Extreme Science and Engineering Discovery Environment (XSEDE), which is supported by National Science Foundation grant number ACI-1053575, and the North Carolina State University High Performance Computing Center.

---

Figure S1. Power (Y-axis) and minor allele frequency (X-axis) of single-stage association tests for three different genetic architectures and three levels of data imbalance. Curves represent loess estimates of the mean and 95% confidence interval of power at each minor allele frequency.

Figure S2. Mean squared error of QTL effect estimates for different GWAS methods under three different genetic architectures and three levels of data imbalance.

Table S1 Power of six different GWAS methods to detect causal variants under three different genetic architectures and three levels of data imbalance.

Table S2 False discovery rate of six different GWAS methods under three different genetic architectures and three levels of data imbalance. Two different thresholds for declaring false positives are reported.

Table S3 Bias and mean squared errors of causal loci effect estimates for six different GWAS methods under three different genetic architectures and three levels of data imbalance.

